# Reward network state dynamics track ASD symptom severity but not Diagnosis

**DOI:** 10.64898/2026.06.12.731591

**Authors:** Jannis Breßgott, József Arató

## Abstract

Autism spectrum disorder (ASD) is a heterogeneous developmental condition characterized by repetitive behaviors and social communication difficulties. The reward network has been specifically implicated in the social deficits associated with ASD for several decades. While modern neuroimaging techniques enable investigation of this cortical-subcortical network, task-based fMRI studies have yielded inconsistent findings, largely due to insufficient sample sizes and heterogeneous reward paradigms. More recently the brain at rest has been leveraged to examine reward network connectivity in large-scale datasets. While previous studies have identified associations between network organization, diagnostic group, and individual-level clinical variables, these findings have largely remained at the trend level. However, prior research has focused exclusively on static connectivity patterns, neglecting the temporal dynamics inherent in brain activity.

In this study, we analyzed a large multi-site fMRI dataset (ABIDE I) to examine reward network dynamics in individuals with ASD, with particular emphasis on individual-level variability and its relationship to clinical phenotypes. Following an initial assessment of static connectivity, we employed a Hidden Markov Model (HMM) as our primary analysis method, alongside a sliding window approach with subsequent clustering as a secondary validation step, to characterize the temporal properties of reward network activity. We observed a consistent association between greater occupancy of the most sparsely connected network state and milder verbal communication symptoms, as measured by the Autism Diagnostic Interview-Revised (ADI-R), across both methods. Notably, traditional group-level comparisons between ASD and control groups revealed limited differences, underscoring the importance of individual-level characterization in this heterogeneous condition. These findings demonstrate that temporal dynamics of reward network connectivity capture clinically meaningful variation in ASD beyond static connectivity measures, supporting the value of dynamic approaches for understanding neurodevelopmental disorders.

## 1 Introduction

Autism Spectrum Disorder (ASD) is a heterogeneous neurodevelopmental condition characterized by persistent deficits in social communication and interaction, alongside restricted, repetitive patterns of behavior (WHO, 2019). With a global prevalence of approximately 1 in 100 individuals, ASD imposes substantial burdens across the lifespan (Zeidan et al., 2022). Among the theoretical frameworks proposed to explain its core social difficulties, the Social Motivation Hypothesis (SMH; Chevallier et al., 2012) posits that reduced reward value assigned to social stimuli drives diminished social engagement and the downstream failure to acquire social skills. This directly implicates the brain’s reward network as a key neural substrate of ASD symptomatology. This distributed cortical-subcortical circuit encompasses, but is not limited to, the nucleus accumbens (NAc), dorsal striatum (caudate and putamen), amygdala, anterior cingulate cortex (ACC), ventromedial prefrontal cortex (vmPFC), orbitofrontal cortex (OFC), and insula (Clements et al., 2018; Haber & Knutson, 2010). Importantly, subsequent work has challenged the specificity of the SMH to social stimuli, instead arguing for a broader deficit in reward sensitivity in ASD (Bottini, 2018; Clements et al., 2018; Janouschek et al., 2021), underscoring the importance of a complete characterization of the network and its relationship to the disorder.

Task-based fMRI studies have consistently reported hypoactivation of reward circuitry in ASD, particularly within the striatum and NAc (Dichter, 2012; Kohls et al., 2013), a finding later confirmed by a meta-analysis showing consistent right-hemispheric striatal hypoactivation (Janouschek et al., 2021). However, synthesizing these results has proven difficult due to heterogeneous reward paradigms, small sample sizes, and conflicting findings across studies (Clements et al., 2018; Janouschek et al., 2021). Resting-state fMRI (rs-fMRI) offers key methodological advantages to address these limitations; its standardized acquisition protocol is compatible with clinical and developmental populations, and its simplicity has enabled the assembly of large multi-site datasets such as the Autism Brain Imaging Data Exchange I (ABIDE I; Di Martino et al., 2014). More generally, rs-fMRI has proven valuable for clinical applications and large-scale neuroimaging due to its robustness and relatively low participant burden (Fox, 2010; Van Dijk et al., 2012). In the absence of an imposed task, functional connectivity (FC), the temporal dependency between activation time series of spatially distant regions (Friston, 1994), has become the primary tool for investigating network activity at scale.

Static FC (sFC) studies, which estimate a single connectivity value per region pair across the entire scan (Lurie et al., 2020), have reported alterations within parts of the reward network in ASD, including decreased NAc–ACC connectivity (Padmanabhan et al., 2013; Polk & Ikuta, 2022), though conflicting results persist (Delmonte et al., 2012; Fishman et al., 2018). The most comprehensive within-network sFC investigation to date by Yang et al. (2024) found reduced left ACC–left NAc and left ACC–right amygdala connectivity in ASD, however the associations of these connections with the Social Responsiveness Scale (SRS) and the Autism Diagnostic Observation Schedule (ADOS) scores, respectively, were significant only at uncorrected thresholds. This pattern of contradictory findings and trend-level associations might reflect the statistical challenge posed by ASD heterogeneity, where variation in age, cognitive ability, symptom profile, and co-occurring conditions attenuates group-level effects and brain–behavior relationships (Hull et al., 2017; Lenroot & Yeung, 2013; Lord et al., 2020). Additionally, all prior reward network studies have relied exclusively on static connectivity, treating brain organization as invariant across the scanning session.

This assumption, however, is contradicted by evidence that functional interactions between brain regions fluctuate meaningfully on the order of seconds to minutes (Chang & Glover, 2010). Dynamic functional connectivity (dFC) captures these time-varying properties and has been shown to reveal clinically relevant information beyond sFC, including altered brain state dynamics in ASD such as increased time in hyperconnected states and reduced network flexibility (Li et al., 2020; Qian et al., 2024), as well as alterations in specific regions (Gao et al., 2022; Zhu et al., 2023). However, to our knowledge, no study has yet examined dFC within the reward network in ASD. Given the substantial empirical literature implicating the reward network in ASD symptomatology, a within-network dFC analysis is highly theoretically motivated. Methodologically, this targeted approach also carries a substantially smaller multiple-comparison burden than whole-brain analyses, following the precedent set by within-network investigations in other clinical populations (Du et al., 2016; Sendi et al., 2021a, 2021b).

However, the field lacks an agreed-upon methodological *gold standard* for dFC analysis (Torabi et al., 2024). The prevalent approach over the last decade has been the sliding-window method combined with subsequent clustering (SWC), which has not been without critique (Laumann et al., 2024). Despite these critiques, within-network dFC studies have predominantly continued to rely on this framework (Bonkhoff et al., 2021; Du et al., 2016; Sendi et al., 2021a, 2021b). Among others, a more recent approach for dFC analysis, the Hidden Markov Model (HMM), offers a more principled framework relying on fewer arbitrary parameter choices and has seen a recent increase in interest (Lurie et al., 2020; Vidaurre et al., 2017). However, approximately 80% of published dFC studies between 2022 and 2023 still relied solely on sliding-window approaches (Laumann et al., 2024), leaving findings potentially contingent on a single method. This is concerning given that these methods diverge substantially in their underlying assumptions. Prior work has shown that spatial and temporal agreement between the outputs of two of the most common methods applied to the same data can be as low as 65% (Torabi et al., 2024). Recent work has therefore called for the application of two or more methods to achieve more robust results (Torabi et al., 2024).

To address these gaps, we analyzed a large-scale fMRI dataset to characterize reward network dFC in individuals with ASD and typically developing controls, employing the HMM as the primary analysis method, with SWC serving as a secondary validation step in a dual-method design intended to yield highly robust findings. Following site harmonization, we first assessed static connectivity, allowing us to evaluate the added value of the dynamic approach. Afterwards, we extracted dynamic network metrics and examined their associations with ASD diagnostic status and dimensional clinical measures.

## 2 Methods

### 2.1 Subjects

Data were drawn from the Autism Brain Imaging Data Exchange I (ABIDE I), an openly shared multi-site neuroimaging dataset comprising resting-state fMRI and phenotypic data from individuals with ASD and typically developing controls (TD) (Di Martino et al., 2014). Quality-based exclusion criteria were applied, with particular attention to the demands of dFC analysis, in addition to standard criteria such as excessive head motion (mean framewise displacement > 0.5 mm). Given that the present study pursued two distinct analytical objectives, group-level ASD versus TD comparisons and dimensional brain–behavior associations with clinical phenotypes, two separate samples were defined. For group comparisons, a sample of *N* = 430 participants was retained (ASD: *n* = 215, TD: *n* = 215), while clinical association analyses were restricted to an ASD-only sample (*N* = 246) with complete entries on relevant clinical measures. Both samples were restricted to male participants with full-scale IQ ≥ 70. To account for systematic inter-site variability arising from heterogeneous scanner hardware and acquisition protocols, ComBat harmonization, as implemented in the NeuroCombat toolbox (Fortin et al., 2018), was applied prior to all analyses, preserving biological variance associated with diagnostic group and the clinical covariates of interest.

### 2.2 Preprocessing and ROI Definition

Preprocessing followed the Configurable Pipeline for the Analysis of Connectomes ([CPAC](https://fcp-indi.github.com/)) pipeline, including slice-timing correction, motion realignment, spatial normalization to MNI space, and bandpass temporal filtering (0.01–0.1 Hz). All included scans had a repetition time (TR) of 2 s, ensuring temporal consistency across participants. Critically, global signal regression (GSR) was applied to reduce the influence of non-neural confounds, as recommended in the analysis of dFC and brain–behavior relationships (Ahrends & Vidaurre, 2023; Li et al., 2019), though we acknowledge that GSR remains debated in the FC literature (Murphy & Fox, 2017). To assess the sensitivity of our findings to this preprocessing choice, all primary brain–behavior and group-comparison analyses were repeated on a non-GSR-preprocessed version of the data (see Appendix 5.3). Reward network nodes were defined following the coordinates reported by Yang et al. (2024), drawn from prior meta-analytic and empirical literature (see Appendix Table 1). For each participant, BOLD time series were extracted by averaging the signal across all voxels within each 6 mm ROI sphere in accordance with prior work (Yang et al., 2024), yielding a regional time series matrix used for all subsequent connectivity analyses.

### 2.3 Analysis

As a reference baseline, static functional connectivity (sFC) was estimated by computing pairwise Pearson correlations between all ROI time series across the full scan duration, yielding a single symmetric connectivity matrix per participant and a group-specific average matrix. Fisher’s *r*-to-*z* transformation was applied prior to statistical testing. Group differences (ASD vs. TD) and associations between connectivity values and clinical measures were assessed using general linear models, with age, site, mean framewise displacement, and full-scale IQ (FIQ) included as covariates.

To assess dFC, the Hidden Markov Model was employed as the primary analytical method. Alongside this, sliding window with subsequent clustering (SWC) was utilised to ensure that the results were robust and not strictly contingent on the assumptions of a single methodological approach.

#### 2.3.1 Hidden Markov Model

The HMM was applied directly to the preprocessed ROI time series without prior windowing, as introduced to fMRI analysis by Vidaurre et al. (2017) and implemented via the `glhmm` Python package (Vidaurre et al., 2025). Under this formulation, the observed fMRI signal at each time point *t* is modeled as arising from one of *K* latent states, governed by initial state probabilities ***π***,a *K × K* transition matrix **A**, and state-specific emission parameters. Each state *k* emits observations according to a multivariate Gaussian distribution:

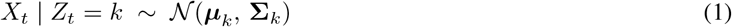

where ***µ***_*k*_ and **Σ**_*k*_ are the state-specific mean vector and covariance matrix, respectively. All model parameters were jointly estimated via variational Bayes inference (Vidaurre et al., 2017, 2025). State-specific functional connectivity was characterised by Pearson correlations obtained by normalising **Σ**_*k*_ to the corresponding correlation matrix.

#### 2.3.2 Sliding Window with K-Means Clustering

A temporal sliding window was applied to each participant’s ROI time series using a Gaussian-tapered window (*ω* = 6) to reduce boundary artifacts and autocorrelation introduced by overlapping windows (Allen et al., 2014). The main analysis was conducted with a window size of 20 time points (40 seconds) and a sliding length of 1 time point (2 seconds). The resulting set of windowed connectivity matrices across all subjects was then submitted to *k*-means clustering, yielding *K* discrete brain states represented as cluster centroids. Since a known weakness of the approach is sensitivity to arbitrary window-length selection, an agnostic multi-window strategy was employed, evaluating several window lengths within the empirically recommended range of 20–60 seconds (Lurie et al., 2020), and the consistency of results across window sizes was assessed (see Appendix).

#### 2.3.3 Model Evaluation & State Selection

State selection was primarily guided by the HMM, which offers a principled measure of model fit in the form of Free Energy (FE). Compared to classical maximum-likelihood approaches, FE has been proposed as a more appropriate criterion for model fit in HMMs applied to fMRI data, as it balances model accuracy against complexity through a regularisation term that penalises overfitting. It has subsequently also been proposed as a selection criterion for the number of states (Rezek & Roberts, 2005; Vidaurre et al., 2025).

To fit the model and to facilitate state selection, we initialised the HMM for each *K* across 100 random seeds and trained each initialisation for 100 iterations, following the Best-Ranked HMM practice (Alonso & Vidaurre, 2023). However, consistent with prior reports (Lin et al., 2022), FE decreased monotonically without a global minimum, though displaying a relative deceleration in the rate of decrease around *K* = 6, whereas model entropy increased across all *K* for both methods (Figure 1). This limitation has been acknowledged in the literature, with the recommendation that state selection should not rest on quantitative criteria alone but also take biologically plausible levels of complexity into account (Vidaurre et al., 2025). *K* = 6 was selected on the basis of the subtle elbow in FE, in line with prior single-network work (Sendi et al., 2021a), and balancing dynamic richness against interpretability. The same value was adopted for the SWC analysis to ensure methodological consistency and enable direct comparison between the two approaches. Acknowledging the inherently subjective nature of this choice, the primary analyses were repeated with *K* = {3, 4, 5, 7}, spanning the range of *K* employed in comparable within-network dFC studies (Du et al., 2016; Sendi et al., 2021b) (see Appendix 5.2).

**Figure 1:**
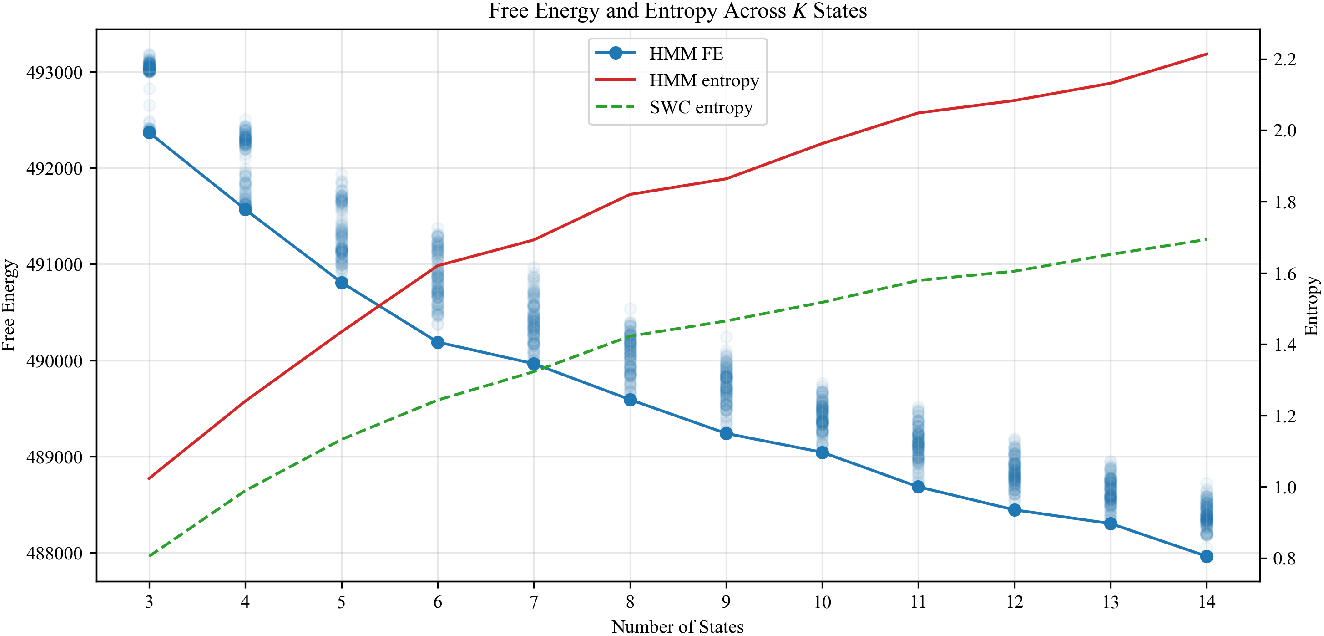
Free Energy and Entropy for models across *K* states. Light blue points show 100 FE initialisations per state.

#### 2.3.4 State Metrics and Statistical Analysis

For each method, six states were derived and subsequently reordered to facilitate comparison based on Euclidean distance using the Hungarian algorithm (Kuhn, 1955). Alongside state-specific metrics such as transition probabilities, the following temporal summary metrics were computed for each subject: fractional occupancy (the proportion of total scan time spent in a given state), mean lifetime (the average consecutive dwell time), and switching rate (the number of state transitions per unit time). Group differences (ASD vs. TD) were tested using permutation-based *t*-tests. Static FC brain–behavior associations were assessed using permutation-based correlation tests. Dynamic associations between state metrics and clinical scores were evaluated using the permutation-based framework introduced by Larsen et al. (2026), in which multivariate associations across state occupancies were assessed using *F* -regression and univariate associations using *t*-tests.

## 3 Results

### 3.1 Static FC Analysis

Static FC analysis across all reward network ROI pairs yielded no significant group differences between ASD and TD participants after FDR correction. At the uncorrected level, 9 connections showed trend-level differences (*p<* 0.05 uncorrected), most prominently involving connectivity between the left ACC and left NAc, and between the vmPFC and left NAc, consistent with the direction of effects reported in prior rs-fMRI studies of the reward network in ASD (Padmanabhan et al., 2013; Polk & Ikuta, 2022; Yang et al., 2024).

Dimensional associations between static reward network connectivity and clinical phenotype measures were examined, with a total of 23 ROI pairs showing uncorrected associations (*p<* 0.05). None of these associations survived FDR correction (all *p*_FDR_ *>* 0.81), confirming the pattern of trend-level, uncorrected effects characteristic of prior static FC investigations of the reward network in ASD (Yang et al., 2024). As anticipated, a single time-averaged summary statistic does not reliably capture clinically meaningful variation in this heterogeneous condition, reinforcing the rationale for a dynamic analysis as the primary approach.

### 3.2 Dynamic FC Analysis

#### 3.2.1 Group Comparison

States were derived on the full sample, comprising equal proportions of ASD and control participants. Permutation-based *t*-tests were then used to compare fractional occupancies, switching rates, and state lifetimes between the two groups for both methods. Consistent with the static FC results, no statistically significant group differences were observed after FDR correction for any temporal metric in either approach. Full results, state characterisations, and associated metrics are reported in Appendix 5.1.

#### 3.2.2 State Characterisation

Subsequent analyses focused on the fully phenotyped ASD-only subsample to examine individual-level brain–behavior associations. Both the HMM and SWC solutions varied in overall connectivity strength, ranging from weakly connected configurations to states characterised by strong positive or negative coupling between specific node pairs. However, the SWC states exhibited stronger differentiation in connectivity magnitude, deviating more markedly from the static baseline than the HMM, which produced comparatively homogeneous states that remained closer to the sFC baseline (HMM *M*_deviation_ = 1.23, *SD* = 0.65; SWC *M*_deviation_ = 2.16, *SD* = 0.57; see Discussion).

#### 3.2.3 Brain–Behavior Associations

Multivariate and univariate permutation tests (50,000 permutations; *F* -regression and Pearson correlation *t*-statistic, respectively) were used to examine associations between the fractional occupancy profiles of the derived states and five clinical measures: ADI-R Social, ADI-R Verbal, ADI-R Restricted and Repetitive Behaviors (RRB), ADOS Social, and ADOS Communication. Age, FIQ, site, and framewise displacement were included as covariates. All *p*-values were corrected using FDR (Benjamini-Hochberg).

For the HMM, the multivariate permutation test identified a significant overall association between the full FO profile and ADI-R Verbal scores (*R*^2^ = .064, *p*_FDR_ = .037). No other clinical measure survived multivariate correction (ADI-R RRB: *p*_FDR_ = .142; all others *p*_FDR_ *>* .499). At the univariate level, the Verbal effect was localised to State 2, where higher fractional occupancy was significantly associated with lower ADI-R Verbal scores (*t* = −3.82, *p*_FDR_ = .010, *r* = −.24). A univariate association of comparable magnitude was also observed between State 2 occupancy and lower ADI-R RRB scores (*t* = −3.30, *p*_FDR_ = .017, *r* = −.21); however, given that the multivariate omnibus test for RRB did not survive correction, this latter effect should be interpreted with caution. No other state–measure pairs survived FDR correction.

For the SWC, no multivariate association survived FDR correction (ADI-R Social: *R*^2^ = .047, *p*_uncorr_ = .040, *p*_FDR_ = .202; all others *p*_FDR_ ≥.328). At the univariate level, however, higher fractional occupancy of SWC State 2 was significantly associated with lower ADI-R Social (*t* = −3.08, *p*_FDR_ = .043, *r* = −.20) and lower ADI-R Verbal scores (*t* = −2.99, *p*_FDR_ = .043, *r* = −.19), with a further trend-level association for ADI-R RRB (*t* = −2.75, *p*_FDR_ = .060). All ADOS subscales remained non-significant in both methods (all *p*_FDR_ *>* .653). The full corrected results are displayed in Figure 3.

Critically, both methods converged on State 2 as the primary correlate of parent-reported symptom severity. Greater time spent in State 2 was consistently associated with lower, that is, less severe, ADI-R Verbal scores across both approaches and method specific findings for the Social and Repetitive Behavior Subscales. ADI-R Verbal emerged as the sole measure reaching corrected significance in both methods, representing the clearest point of convergence. Notably, this convergence should be interpreted with some caution: the multivariate *F* -test, which serves as a more conservative omnibus test less susceptible to inflated Type I error, was only significant for the HMM. The SWC univariate findings are therefore best understood as corroborating rather than independently sufficient evidence. State 2 for both methods is displayed in more detail in Figure 4.

A robustness analysis was conducted to assess the sensitivity of the findings to the choice of the number of states. Specifically, the analyses were repeated for *K* ={3, 4, 5, 7} . The solutions with 3, 4 and 5 states showed patterns that were largely consistent with the 6-state solution across both methods. In particular, ADI-R Verbal remained significant in the multivariate analysis, and the univariate associations preserved their direction and were of comparable magnitude.

For the 7-state solution, the direction of the observed effects remained stable; however, none of the multivariate associations reached statistical significance. This may reflect reduced statistical power or increased model complexity rather than a substantive change in the underlying relationships.

#### 3.2.4 Sensitivity to Global Signal Regression

Given the ongoing debate regarding global signal regression in dFC analysis (Murphy & Fox, 2017), all primary analyses were repeated on a non-GSR-preprocessed version of the data (see Appendix 5.3); for both methods, states were matched to the GSR solutions. The direction of the primary association was preserved: in both the HMM and SWC, the most sparsely connected state (State 2) showed negative correlations with the ADI-R subscales, mirroring the main analysis. However, neither the multivariate nor the univariate effects reached FDR-corrected significance under this pipeline, and the same pattern of non-significant group-level differences was observed. The wider state profiles diverged modestly: in the SWC, State 2 remained the dominant ADI-related state, whereas in the HMM a second state (State 4) showed a positive correlation of comparable magnitude with the ADI-R subscales. Notably, this State 4 corresponds to the second most densely connected configuration in the HMM solution, characterised by broad positive coupling across the reward network; the opposing sign of its association is therefore directionally coherent with the sparse-state finding, in that greater time spent in a strongly coupled configuration tracks more severe symptoms while greater time in the decoupled State 2 tracks milder ones. The directional consistency of the State 2 effect provides modest support for the robustness of the underlying relationship, while the loss of significance indicates that its magnitude is partially contingent on this preprocessing choice, consistent with reports that GSR can strengthen brain–behavior associations in rs-fMRI (Murphy & Fox, 2017).

## 4 Discussion

### 4.1 Static Connectivity as a Baseline

At FDR-corrected thresholds, the present static analysis did not reproduce the within-reward-network findings reported in the largest prior investigation of this network (Yang et al., 2024), namely weaker left ACC–left NAc and left ACC–right amygdala connectivity in ASD. The uncorrected trend toward weaker left ACC–left NAc connectivity in our ASD group is nonetheless directionally consistent with that finding, as were the corresponding brain–behavior correlations, which preserved the same direction at trend level without reaching significance. Given the small effect sizes Yang et al. reported (Cohen’s d between -0.24 and -0.41), our more tightly matched and smaller sample may be underpowered to detect effects of this magnitude after correction, which together suggests a consistent but weaker underlying signal. These sFC results thus provide a baseline against which the added value of the dynamic approach can be evaluated: clinically meaningful variation in reward network organization might not be reliably captured by a single time-averaged summary statistic.

### 4.2 Absence of Group-Level Differences in Dynamic Metrics

Consistent with the static analysis, no significant group differences between ASD and TD participants were observed for any temporal metric—fractional occupancy, state lifetime, or switching rate—across either the HMM or the SWC following FDR correction. This null result is unsurprising given the well-documented challenge of detecting reliable group-level effects in ASD neuroimaging, where substantial within-group heterogeneity in age, cognitive profile, symptom severity, and co-occurring conditions routinely attenuates between-group contrasts (Hull et al., 2017; Lenroot & Yeung, 2013; Lord et al., 2020). Prior dFC studies reporting group differences have generally done so using whole-brain analyses that capture broader variance, whereas the spatial scope of the present study is deliberately restricted to the reward network, an approach that has precedent in focused within-network investigations of other conditions (Du et al., 2016; Sendi et al., 2021a, 2021b). The uncorrected trends present in both methods, most notably slightly lower fractional occupancy in states 2 and 3 for the ASD group in the HMM, could reflect a genuine but weak group signal that is overwhelmed by within-group variance at corrected thresholds. Rather than interpreting these null findings as evidence against reward network dysfunction in ASD, they reinforce the case made by the sFC results: categorical group comparisons are likely insufficient to characterize the heterogeneous neural basis of this condition, and individual-level, dimensional approaches are necessary.

### 4.3 Sparse State Occupancy and Verbal Communication

The primary finding of this study is a significant association between greater fractional occupancy of the most sparsely connected network state and lower, that is, less severe, ADI-R Verbal Communication scores, converging across both dFC methods. For the HMM, this constituted the sole multivariate effect surviving correction and was robustly localised to State 2 at the univariate level, together with a univariate association with the ADI-R RRB score that was significant only in the HMM but not in the SWC. Conversely, the SWC uniquely yielded a corrected univariate association with ADI-R Social. ADI-R Verbal was the sole measure reaching corrected significance in both methods, constituting the clearest point of cross-method convergence. In general, the effect of State 2 is consistently negative across all ADI-R subscales and methods and retains that direction for most of the robustness checks. This convergence is particularly notable because, as discussed in Section 4.4, the two approaches produce states of markedly different spatial sharpness and temporal stability; that both independently implicate the same sparsely connected state as the primary neural correlate of verbal communication severity argues against the association being a methodological artifact.

State 2 was characterised by widespread reductions in reward network coupling relative to the static baseline, with particularly attenuated connectivity between the vmPFC, ACC, amygdala, and insula. This sparse connectivity profile aligns with static reports of reduced ACC–amygdala coupling in ASD, which have been correlated with the severity of social communication deficits (Yang et al., 2024). In addition, reduced connectivity between the insula and the amygdala has been proposed to impair integration of affective and social signals (Von Dem Hagen et al., 2013). The present results extend this picture temporally: it is the proportion of time the network spends in a decoupled configuration, rather than a single average connectivity value, that covaries with individual differences in verbal communication ability. The direction of the effect, with more time in the sparse state predicting better verbal outcomes, may initially appear counterintuitive but is consistent with the view that low-connectivity states might represent a functional reconfiguration period, or a period of out-of-network connection, that has been implicated in associations with cognitive performance and dimensional psychopathology (Fu et al., 2025). Under this interpretation, individuals with more severe verbal communication difficulties may spend less time in this disengaged mode and correspondingly more time locked into atypical coupling configurations, potentially connected with other networks, paralleling reports of increased time in hyperconnected states and reduced network flexibility in broader ASD dFC studies (He et al., 2018; Li et al., 2020). While the question of out-of-network dynamics cannot be directly addressed by this study, we acknowledge prior work that has made progress in investigating whole-brain dynamics (Lin et al., 2022; Qian et al., 2024) and seed-based analyses focusing on specific regions (Gao et al., 2022; Zhu et al., 2023). Nevertheless, we argue that, given the substantial prior work concerning reward network involvement in ASD, it is both statistically principled and theoretically motivated to restrict the analysis to this specific network. Future work, however, should characterise the temporal dynamics of the reward network relative to the rest of the brain to obtain a fuller picture of within- and between-network connectivity (Alonso et al., 2025).

The observed within-network pattern does warrant explicit reconciliation with the Social Motivation Hypothesis, which links reduced reward-network engagement to more severe social-communicative impairment. Our finding, that less within-network coupling accompanies milder verbal impairment, appears to run counter to this prediction. We argue, however, that this result complicates and refines the framework instead of directly contradicting it. First, the directional claims of the SMH rest on the task-evoked valuation of social stimuli, whereas the present analysis quantifies the proportion of time the intrinsic network spends in a decoupled configuration. To our knowledge, there is no specific reason that greater resting within-network synchrony should directly index healthier reward function. Second, the effect was specific to parent-reported verbal communication measures, rather than to all measured social scales, showcasing a relationship with a specific facet instead of a complete inversion. We acknowledge, nonetheless, that if resting within-network coupling is treated as a proxy for reward engagement, the present direction would instead constitute evidence against the social-specific form of the SMH, which then would converge with existing critiques of its specificity (Bottini, 2018; Clements et al., 2018).

The specificity of the association to parent-reported ADI-R scores, rather than the clinician-administered ADOS, is surprising but theoretically interpretable. The ADOS captures observable communicative behaviour during a structured interaction and may reflect situationally bound symptom expression that is less tightly coupled to the trait-level neural organisation indexed by resting-state dynamics (Lord et al., 2020). The ADI-R, by contrast, aggregates caregiver reports of habitual behaviour across developmental history and multiple contexts, which may be more closely aligned with the stable individual differences in reward network temporal dynamics captured here (Lord et al., 2020).

### 4.4 Between Method Considerations

Given their distinct underlying assumptions, the two methods naturally produced differing state profiles, despite showing considerable convergence in capturing brain–behavior relationships. This divergence is immediately evident in a qualitative comparison of the derived brain states (see Figure 2). SWC yielded highly distinct states characterized by pronounced hyper- or hypo-connectivity in specific regions, whereas the states identified by the HMM were comparatively subtle and exhibited greater spatial similarity to one another. This discrepancy stems from the core mechanics of each approach. The SWC method relies on *k*-means clustering, which minimizes within-cluster distances (Hartigan & Wong, 1979). By optimizing for compact clusters, *k*-means tends to produce well-separated centroids as an emergent consequence of assigning points to their nearest center. Conversely, the HMM is a probabilistic temporal model that does not explicitly maximize spatial separation; instead, it optimizes the likelihood of the entire time series to best explain its continuous temporal evolution (Vidaurre et al., 2025). Consequently, while states across the two methods were matched based on spatial similarity (i.e., Euclidean distance) to facilitate comparison, they do not represent identical ground-truth configurations.

**Figure 2:**
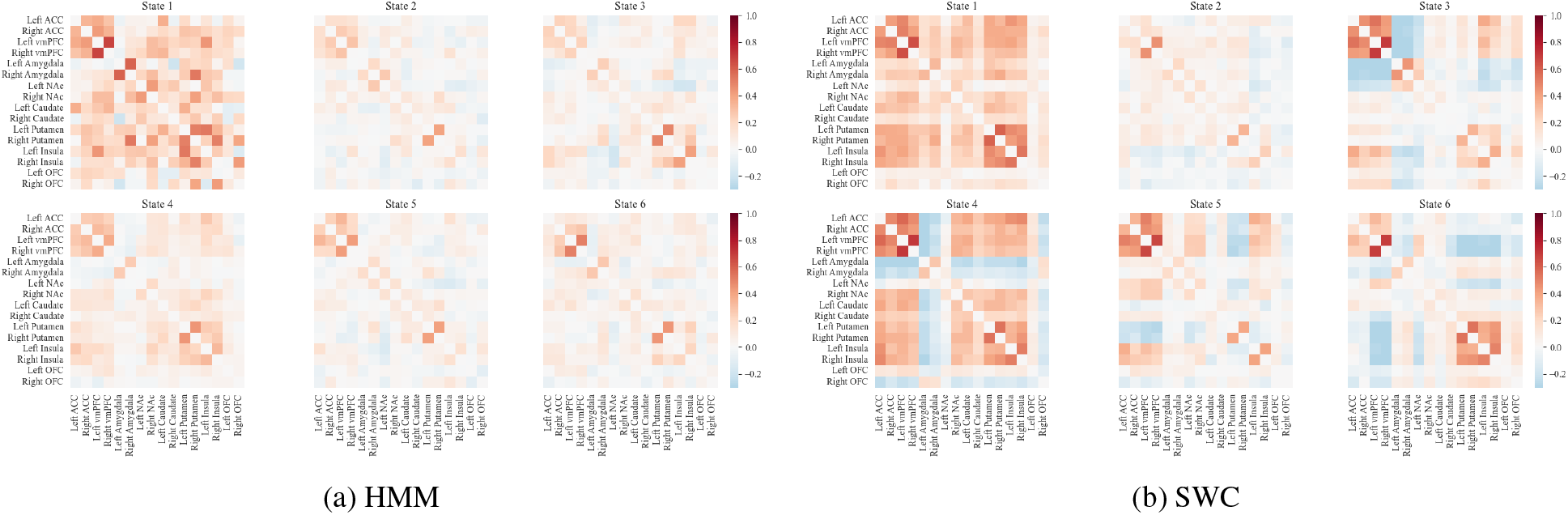
States derived from both methods, reordered by Euclidean distance.

**Figure 3:**
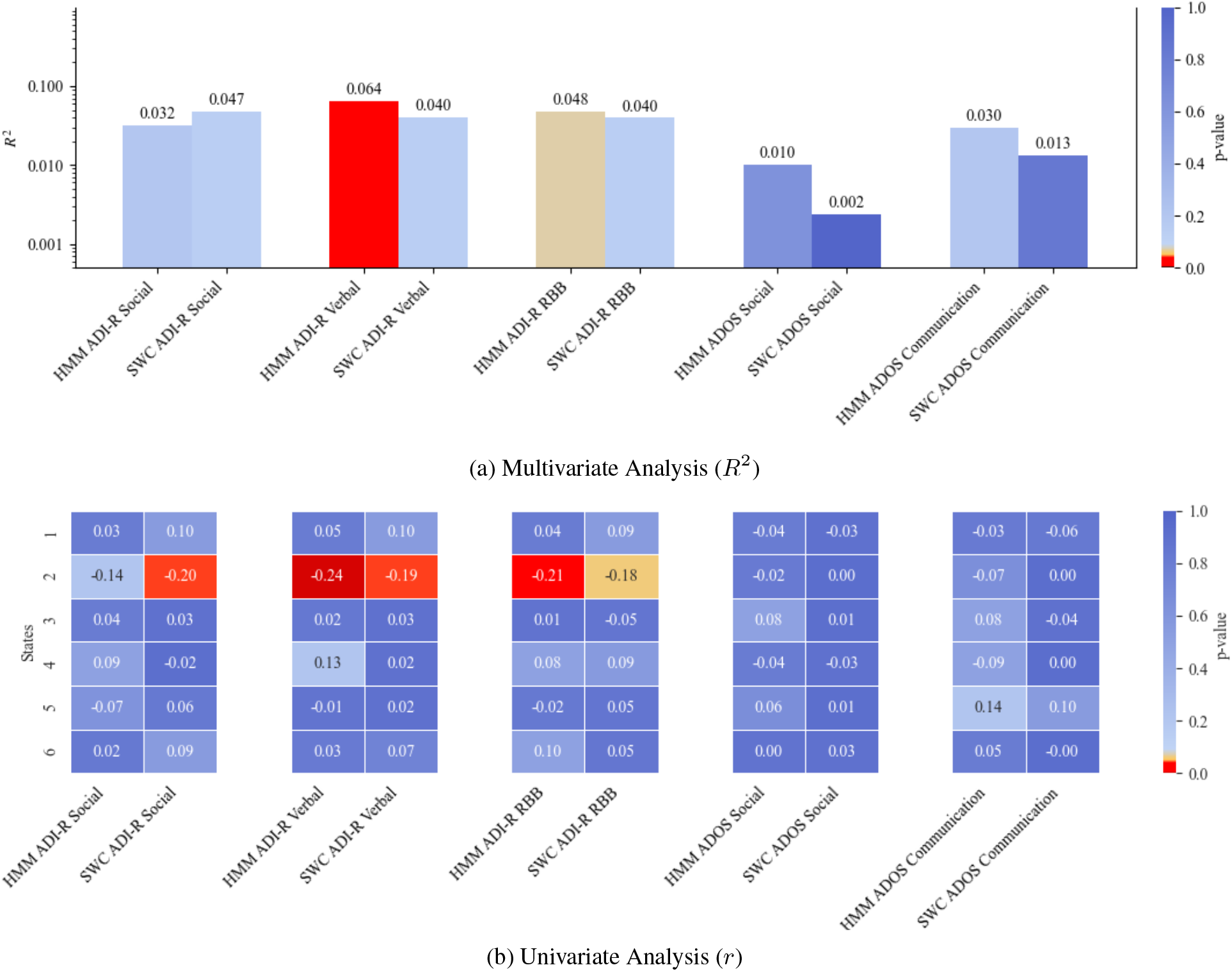
Brain–Behavior Results, Annotations display effect sizes, Color indicates p-values

**Figure 4:**
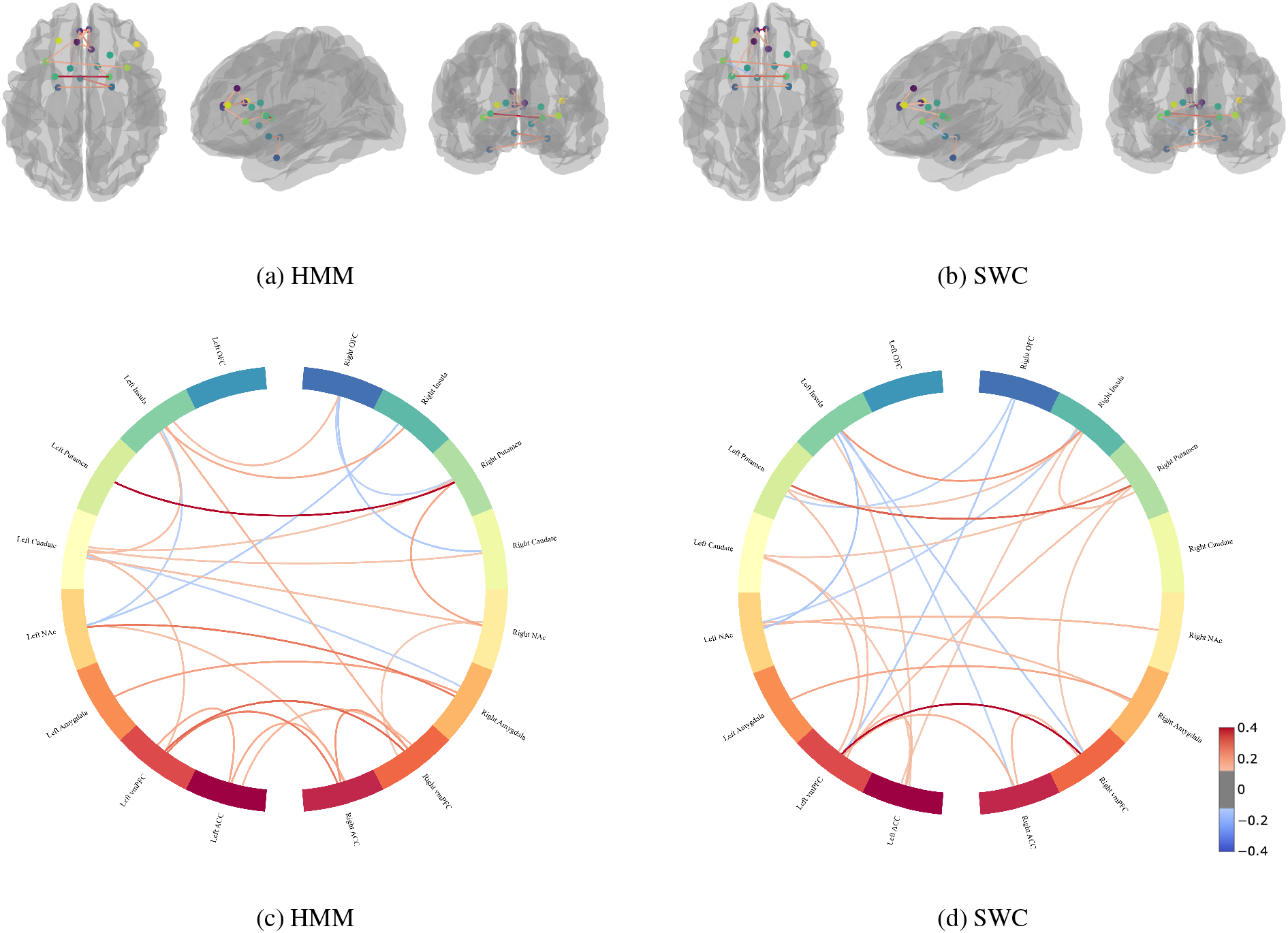
3D Visualization and Connectome plot of State 2 for both Methods at 90% Threshold

These algorithmic differences also manifested in divergent temporal dynamics. The mean lifetime of SWC states was approximately 6 seconds, compared to 8.5 seconds for HMM states. Accordingly, the SWC method yielded a state switching rate nearly one and a half times higher than that of the HMM. Rather than reflecting true biological instability, this elevated switching rate is likely a methodological artifact. SWC relies on independent, hard assignments for each discrete sliding window, making it highly susceptible to noise and minor fluctuations that can trigger spurious transitions. In contrast, the HMM decodes a globally optimal state sequence via the Viterbi algorithm at the temporal resolution of the raw data, where each assignment is jointly constrained by transition probabilities learned across the entire time series. A state switch is therefore only decoded when it is consistent with the global temporal structure of the recording, conferring robustness to local noise. The HMM’s more stable temporal output thus likely reflects this architectural constraint rather than a divergence in the underlying biological signal.

One possible explanation for the improved alignment might be the reduced parameter space of a targeted single-network analysis (i.e., fewer ROIs compared to whole-brain analyses) combined with a robust variational Bayes HMM implementation. Ultimately, these findings validate the dual-method design: rather than yielding contradictory evidence, the use of complementary methodologies improves confidence in the observed reward network dynamics, a robustness that prior single-method within-network clinical studies (Du et al., 2016; Sendi et al., 2021a, 2021b) were unable to provide. Future work should systematically explore how varying sample compositions and parameter choices influence cross-method convergence, thereby informing best practices for multi-method validation in dynamic functional connectivity research.

### 4.5 Limitations

Several limitations qualify these findings. First, despite ComBat harmonization, residual site-related variance from the multi-site ABIDE I design cannot be entirely excluded and may contribute noise to the brain–behavior associations. Second, the present analysis required a fully phenotyped sample, which reduced usable participants to approximately 25% of the total ABIDE I dataset; the sample was further restricted to male participants with FIQ ≥70, limiting generalizability to females, in whom reward network organisation in ASD may differ (Lenroot & Yeung, 2013), and to individuals with intellectual disability. Future research should seek to leverage the additional capacity of complementary large-scale datasets such as ABIDE II or the UK Biobank to increase sample size and test the robustness of the present findings. Third, the choice of *K* = 6 states, though supported by robustness checks across *K* = {3, 4, 5}, rests on a combination of quantitative criteria and biological plausibility rather than an algorithmic optimum; the 7-state solution in particular failed to preserve multivariate significance, indicating that the present findings are sensitive to model complexity. Fourth, the study design is correlational and cross-sectional: it remains unknown whether sparse-state occupancy is a precursor to communication difficulties, a consequence of atypical reward processing, or a correlated epiphenomenon. Fifth, scan durations varied across ABIDE I sites and were truncated to a common length, placing the effective recording at the lower bound recommended for reliable dFC estimation; longer and more uniform recordings would increase the reliability of individual-level dFC estimates and the statistical power of brain–behavior analyses (Laumann et al., 2024). Sixth, the reward network definition followed prior work that, although grounded in meta-analytic evidence, does not include all regions implicated by other accounts of reward circuitry; the spatial scope of the present findings may therefore not fully generalise to alternative network parcellations. Future work should seek to replicate the sparse-state finding in prospective, harmonized cohorts with extended scan protocols, evaluate alternative network definitions, and systematically probe the influence of preprocessing choices such as global signal regression on the stability of the observed associations.

### 4.6 Conclusion

In this work, we investigated reward network dynamics in ASD, motivated by the long-standing question of how reward circuit impairments relate to symptom severity and whether those impairments extend beyond purely social deficits. Acknowledging the absence of an agreed-upon methodological gold standard, we employed two approaches with distinct underlying assumptions to assess dynamic connectivity and contrasted these against a static baseline. Consistent with the within-group heterogeneity inherent in ASD, no group-level differences between ASD and control participants were detected; instead, individual-level associations emerged between reward network temporal dynamics and verbal communication impairments. Notably, both methods converged on a negative association between fractional occupancy of the most sparsely connected network state and ADI-R verbal severity scores, suggesting that greater time spent in a decoupled network configuration is linked to milder verbal impairments. Crucially, this association was not detectable using static connectivity analysis. Future work should build on these results by characterising how sparse within-network activity relates to between-network connectivity, leveraging that fuller picture to clarify the complete role of the reward circuit in the social and communicative phenotype of ASD.

## Data Availability

All data analyzed in this study are openly available through the Autism Brain Imaging Data Exchange I repository (http://fcon_1000.projects.nitrc.org/indi/abide/), as described in Di Martino et al. (2014). Preprocessed neuroimaging data (CPAC pipeline) and phenotypic information were obtained via the Preprocessed Connectomes Project (http://preprocessed-connectomes-project.org/abide/).

## Code Availability

All analysis code will be made publicly available on GitHub upon publication.

## Author Contributions

J.B.: Conceptualization, Methodology, Software, Formal analysis, Investigation, Data curation, Visualization, Writing – original draft, Writing – review & editing. J.A.: Conceptualization, Methodology, Supervision, Project administration, Resources, Writing – review & editing.

