## Supplementary Material for "Reward network state dynamics track ASD symptom severity but not Diagnosis"

### 1 Appendix

Table 1: Montreal Neurological Institute coordinates and sources of regions of interest for the reward system.

| | x | y | z | Source literature |
|---|---|---|---|---|
| Left ACC | -6 | 32 | 28 | (Janouschek et al., 2021) |
| Right ACC | 6 | 28 | 16 | (Janouschek et al., 2021) |
| Left vmPFC | -4 | 44 | -14 | (Janouschek et al., 2021) |
| Right vmPFC | 4 | 46 | -12 | (Janouschek et al., 2021) |
| Left amygdala | -26 | -2 | -26 | (Clements et al., 2018) |
| Right amygdala | 26 | -2 | -16 | (Clements et al., 2018) |
| Left NAc | -4 | 6 | -12 | (Clements et al., 2018) |
| Right NAc | 12 | 16 | -4 | (Clements et al., 2018) |
| Left caudate | -12 | 12 | 16 | (Clements et al., 2018) |
| Right caudate | 22 | 24 | 12 | (Clements et al., 2018) |
| Left putamen | -26 | 6 | 6 | (Clements et al., 2018) |
| Right putamen | 22 | 6 | -2 | (Zürcher et al., 2021) |
| Left insula | -34 | 20 | 2 | (Clements et al., 2018) |
| Right insula | 38 | 14 | -4 | (Clements et al., 2018) |
| Left OFC | -22 | 36 | -14 | (Xiao et al., 2022) |
| Right OFC | 44 | 32 | -18 | (Clements et al., 2018) |

*Note.* ACC, anterior cingulate cortex; vmPFC, ventromedial prefrontal cortex; NAc, nucleus accumbens; OFC, orbitofrontal cortex.

#### 1.1 Group Comparison

For the HMM, the clearest uncorrected trends emerged in fractional occupancy, where the control group showed slightly higher mean occupancies in States 2 and 3 (State 2: $\Delta M = -.012$, $p_{\text{uncorr}} = .007$, $p_{\text{FDR}} = .103$, ASD $M = .161$, Control $M = .173$; State 3: $\Delta M = -.009$, $p_{\text{uncorr}} = .022$, $p_{\text{FDR}} = .162$, ASD $M = .167$, Control $M = .175$). A marginal uncorrected difference in the opposite direction appeared in State 1, with slightly higher occupancy in the ASD group ($\Delta M = .009$, $p_{\text{uncorr}} = .046$, $p_{\text{FDR}} = .226$, ASD $M = .157$, Control $M = .148$). All remaining states showed negligible group differences in occupancy. State lifetimes were similarly overlapping, with State 2 showing the strongest (but still non-significant) trend toward longer dwell times in controls ($\Delta M = -.186$, $p_{\text{uncorr}} = .070$, $p_{\text{FDR}} = 1.00$, ASD $M = 3.72$, Control $M = 3.91$); no other state approached uncorrected significance (all $p_{\text{uncorr}} > .24$). Switching rates were nearly identical across all six states (all group means $\approx .107$–$.109$, all $p_{\text{uncorr}} > .14$).

For the SWC, group differences were similarly muted across all metrics, though the strongest trends shifted from occupancy to switching rate. No state approached significance in fractional occupancy after correction (all $p_{\text{FDR}} = 1.00$); State 5 again exhibited the highest overall fractional occupancy across both groups (ASD $M = .315$, Control $M = .320$), suggesting it represents the most frequently visited connectivity configuration in the SWC solution. Marginal uncorrected trends emerged in switching rate, where the ASD group showed higher rates in State 4 ($\Delta M = .031$, $p_{\text{uncorr}} = .046$, $p_{\text{FDR}} = .660$, ASD $M = .217$, Control $M = .187$) and controls showed higher rates in State 1 ($\Delta M = -.030$, $p_{\text{uncorr}} = .090$, $p_{\text{FDR}} = .660$, ASD $M = .202$, Control $M = .232$). Lifetime in State 1 also showed a non-significant trend toward longer dwell times in the ASD group ($\Delta M = .468$, $p_{\text{uncorr}} = .105$, $p_{\text{FDR}} = 1.00$, ASD $M = 5.13$, Control $M = 4.66$); no other state approached uncorrected significance. After FDR correction, no test in either model reached the conventional $\alpha = .05$ threshold, paralleling the absence of robust group-level dynamic differences and providing the context for the behavioral association analyses that follow.

#### 1.2 Brain Behavior Robustness

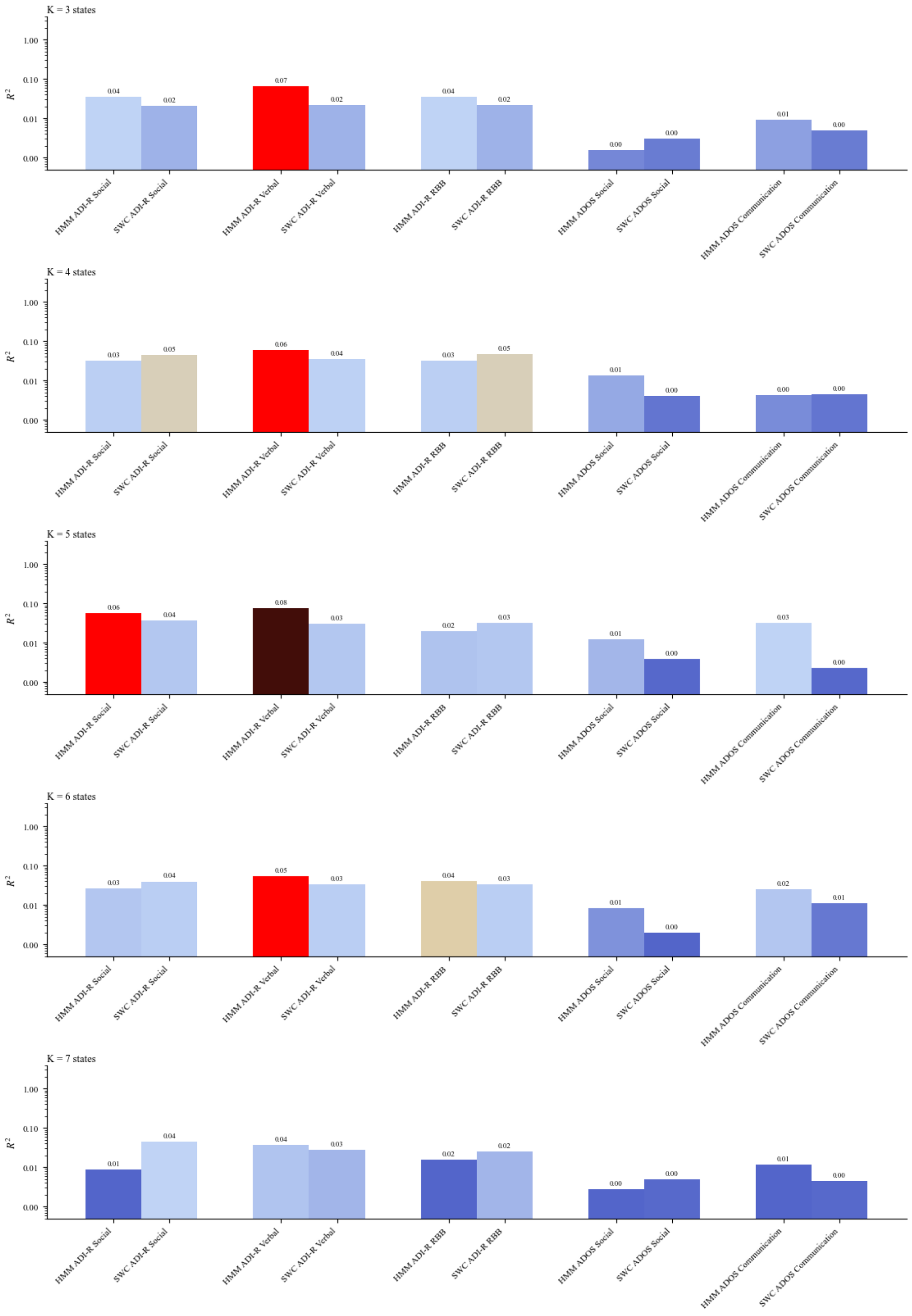


Figure 1: Multivariate Robustness Checks

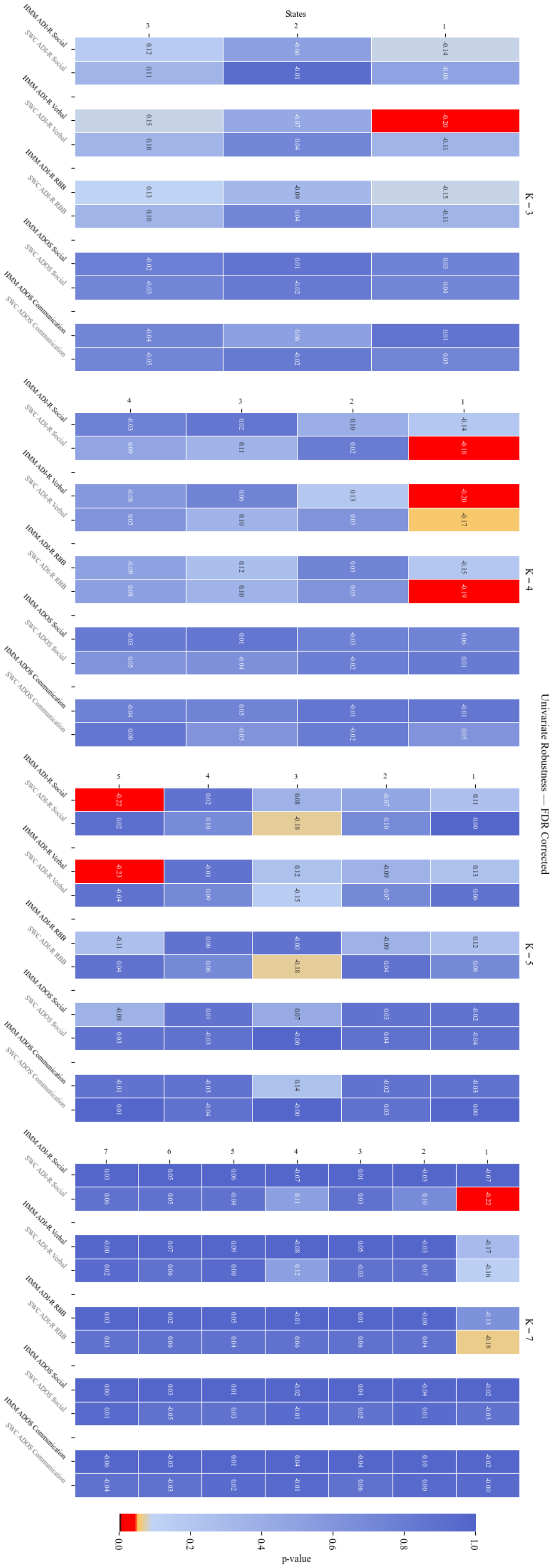


Figure 2: Univariate Robustness Checks

#### 1.3 Non-GSR Results

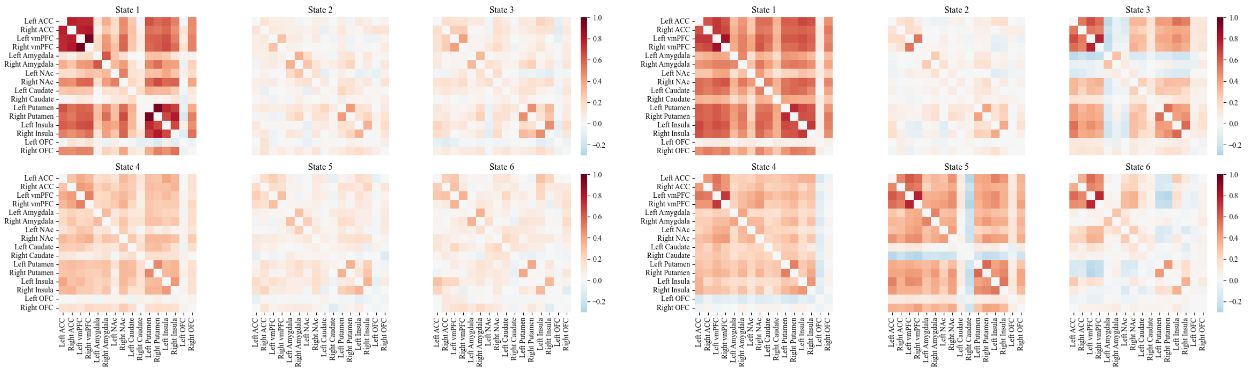

(a) HMM (b) SWC

Figure 3: States derived from both methods for the Non-GSR Data

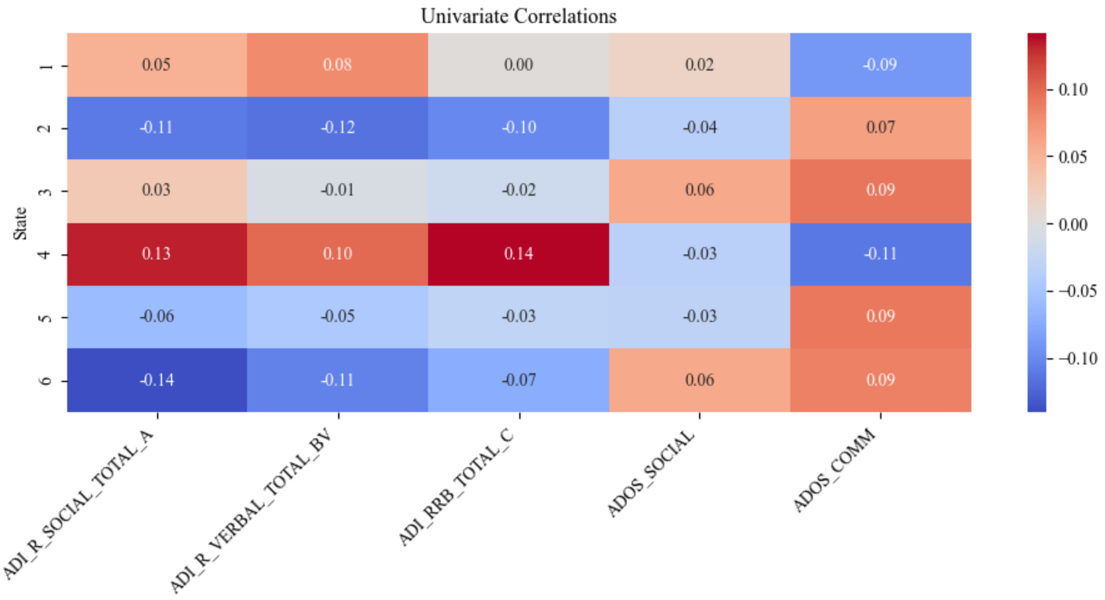


Figure 4: HMM Non-GSR Correlations

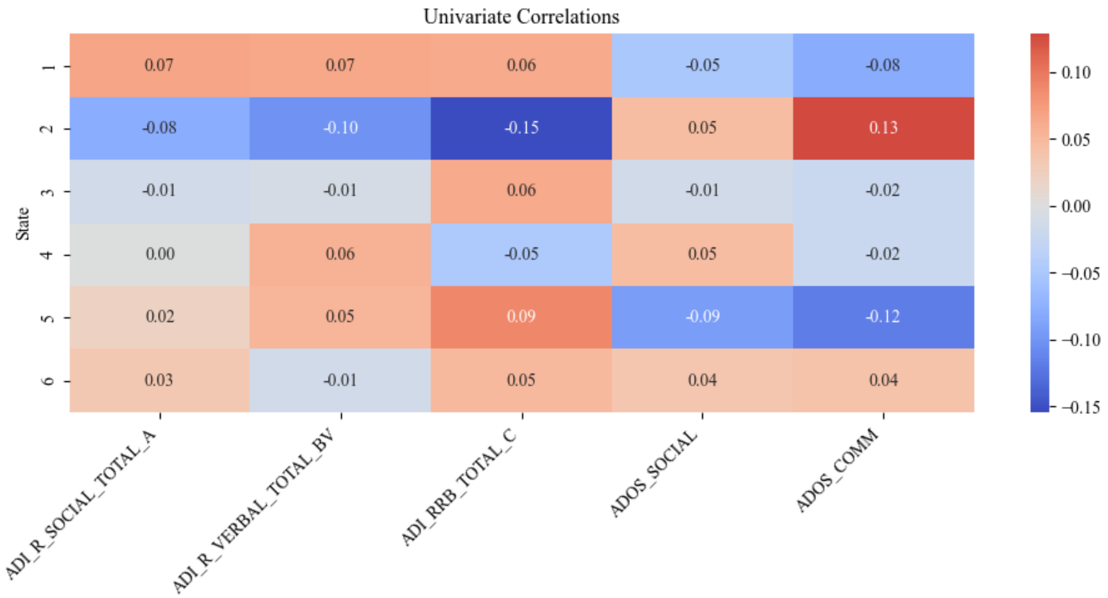


Figure 5: Sliding Window Non-GSR Correlations
